# MicroRNA profiling identifies novel regulators of stem cell function in the adult *Drosophila* intestine

**DOI:** 10.1101/2024.12.30.630748

**Authors:** Perinthottathil Sreejith, Joshuah Yon, Kalina Lapenta, Benoit Biteau

**Affiliations:** Department of Biomedical Genetics, University of Rochester Medical Center, 601 Elmwood Avenue, Rochester, New York USA 14642

**Author notes:** Current address: Department of Biology, Easter New Mexico University, Station 33, 15 S Ave K, Portales NM 88101.

**Keywords:** microRNAs, Drosophila, intestinal stem cells, miR-31a, miR-34

## Abstract

Precise control of stem cell activity is critical to maintain homeostasis and regenerative capacity of adult tissues and limit proliferative syndromes. Hence, stem cell-specific complex regulatory networks exist to exquisitely maintain gene expression and adapt it to tissue demand, controlling self-renewal, fate commitment and differentiation of developing and adults cell lineages. One of the essential and conserved regulatory components that fine-tune gene expression are microRNAs, which post-transcriptionally regulate stability and translation of messengers. microRNAs have been identified as critical stem cell regulators across stem cell populations and organisms. Here, we report the profiling of microRNAs expressed in stem cells and their immediate daughter cells in the *Drosophila* adult intestine. Our analysis identifies over 60 miRs that can be reliably detected in these sorted progenitor cells; a few of these have been reported to control fly intestinal stem cells, but most have yet to be investigated in the adult intestinal lineage. To validate the relevance of our unbiased analysis, we chose to characterize the phenotypes associated with genetic manipulations of two of these microRNAs, miR-31a and miR-34, which are conserved in other organisms, but whose function has not been investigated in the Drosophila midgut. We found that miR-31a acts as anti-proliferation factor and is important for the re-entry of ISC into quiescence after tissue damage. Additionally, we demonstrate that miR-34 is essential for ISC proliferation, but its over-expression also prevents proliferation, highlighting the complexity of miR-mediated control of stem cell function. Altogether, our work establishes a new critical resource to investigate the detailed mechanisms that control stem cell proliferation and intestinal differentiation under homeostatic conditions, in response to tissue damage, or during epithelial transformation and aging.

## Introduction

Many of the major cellular functions such as proliferation, differentiation, growth, and metabolism are regulated by microRNAs (miRs) (Bartel 2004, Jonas and Izaurralde 2015). Though their cellular function appears to vary in eukaryotes, the fact that many microRNA sequences are conserved between distantly related organisms indicates that microRNAs are involved in essential cellular functions and their regulation (Bartel 2004). In *Drosophila*, microRNAs play a crucial role in stem cell biology. For instance, several microRNAs are necessary for the development of Germline Stem Cells (GSCs) during embryogenesis, for gametogenesis and for the regulation of GSC function in adults both in males and females. MicroRNAs are required in the adult ovary in both GSCs and Somatic Stem cells, where they regulate self-renewal and differentiation (Shcherbata 2019). Among the conserved miRs that regulate stem cell function across organisms, the microRNA *let-7* is probably the most studied and targets several niche-specific mRNAs in the fly testis, its down-regulation resulting in age-related loss of GSCs (Toledano, D’Alterio et al. 2012). Another example highlighting the critical role of microRNAs in stem cells across tissues is the miR *bantam*. While it regulates the fate of GSC in the ovary (Yang, Xu et al. 2009), neuronal stem cell fate in the brain (Wu, Lee et al. 2017) and cell growth in the hematopoietic progenitor niche (Lam, Tokusumi et al. 2014), *bantam* is also essential for adult intestinal stem cells (ISCs) self-renewal in the midgut (Huang, Li et al. 2014).

The *Drosophila* adult intestine is maintained by a resident ISC population, which are the only mitotic cells in this tissue. Their division gives rise to daughter cells enteroblasts (EBs) that further differentiate into enterocytes (ECS), or enteroendocrine (EE)-committed daughter cells that give rise to secretory cells (Jiang, Tian et al. 2016). Importantly, ISC proliferation is essential to maintain long term epithelial homeostasis and can be stimulated to promote tissue repair in response to acute damage (Jiang, Tian et al. 2016).

The function of several other miRs has started to be investigated in the *Drosophila* adult intestine. These studies include reports showing that (i) *miR-8* regulates the differentiation of stem cell progenitors in the intestine (Antonello, Reiff et al. 2015); (ii) *miR-305* regulates insulin signaling in ISCs during adaptive homeostasis (Foronda, Weng et al. 2014); (iii) *bantam* controls ISCs self-renewal by regulating Hippo signaling (Huang, Li et al. 2014); (iv) *miR-277* antagonizes fatty acid oxidation in ISCs and is essential for their survival (Zipper, Batchu et al. 2022). While these studies focus on the role of miR in stem cells and progenitors, others have profiled miRs expressed in the intestinal epithelium and non-cell autonomous effect of other miRs on ISCs. *miR-263a* maintains ISC homeostasis by regulating of ENaC (an epithelial sodium channel) which is required for osmotic homeostasis in the midgut epithelium (Kim, Hung et al. 2017); a tissue specific microRNA profiling of the intestinal epithelium identified the role of *miR-958* in the regulation of ISC numbers non-cell autonomously (Mukherjee, Paricio et al. 2021); other small RNA sequencing in the fly intestine identified the conserved role of miR-7 in controlling ISC proliferation (Singh, Hung et al. 2020).

This unbiased approach highlighted the need to better identify miRs expressed in stem and progenitor cells themselves, as these likely control ISC cell-autonomously. However, we are still lacking a comprehensive view of miRs expression, function and regulation in this tissue. To identify the microRNAs with a potential role in the regulation of the stem cell progenitors, we performed a small RNA sequencing of isolated esg-positive cells (Dutta, Buchon et al. 2015). We identified 58 miRs consistently detectable in stem cell and enteroblasts, including miRs with a previously described role in the intestinal lineage. Of these 58 miRs identified in progenitors, we found that *miR-31a* negatively regulates ISC proliferation and *miR-34* is required for ISC cell proliferation. Our results identify progenitor specific miRNAs which are candidates for future functional studies of the role of miRNA-mediated post-transcriptional regulation in ISC self-renewal, fate decision and differentiation.

## Results and Discussion

### Dicer-1 is required in intestinal progenitors to maintain self-renewal capacity

To explore the genetic requirement of the microRNA pathway in intestinal homeostasis, we first knocked-down the expression of Dicer-1 (Dcr-1), a conserved essential component of the microRNA biosynthesis pathway, in intestinal progenitors. To knock-down Dcr-1 specifically in adult ISC/EBs, we used 2 different dsRNA constructs directed against separate regions of the Dicer1 messenger and the esgGal4,UAS-GFP;tubGal80^ts^ (esgGFP^ts^) driver. We found that this genetic perturbation is sufficient to cause significant loss of GFP positive cells in the intestinal epithelium (Figure S1A and S1B). ISCs and EBs can be distinguished by staining against the Notch ligand Delta: ISCs are esg+/Dl+ and EBs are esg+/Dl-. Quantification of Dl+ cells demonstrates that the number of ISCs isn’t significantly different between controls and esgGFP^ts^>dcr-1^RNAi^ expressing animals (Figure S1C). This demonstrates that EBs are largely depleted from the midgut following dcr-1 knock down and that disrupting microRNA biogenesis strongly impairs ISC proliferative capacity. This also suggests that manipulating Dcr-1 is a viable strategy to influence miR levels and allow distinguishing between ISC-and EB-enriched microRNAs.

### Identification of microRNAs expressed in intestinal stem cells and early progenitors

To gain a comprehensive view of the potential miRs involved in the regulation of the ISC lineage in the adult *Drosophila* midgut, we performed small RNA sequencing of isolated esg-positive cells (Figure S2A and S2B). To maximize the opportunity of identifying ISC/EBs-specific specific miRs, around 50,000 GFP-positive cells were FACS sorted in triplicates for each genotype (controls and two Dcr-1 knock-downs) (Figure S2C) and small RNA populations sequenced. Of the 467 miRs annotated in the *Drosophila* genome at the time of these studies (Lu, Baras et al. 2018), 63 were reproducibly detected in at least one of the three genetic conditions, and 38 miRs were detected in all 9 sequenced libraries (Figure 1A and 1B). Specifically, in control cells, 58 miRs presented at least 5 normalized counts in each biological replicate. Using the same criteria, we reliably identified 56 and 41 miRs in cells expressing the #42901 and #34826 UAS-dicer1RNAi constructs respectively. Of these, most microRNAs showed an average count >10 rpkm in the biological triplicates and are presented in a rank order in Figure 1A.

**Figure 1:**
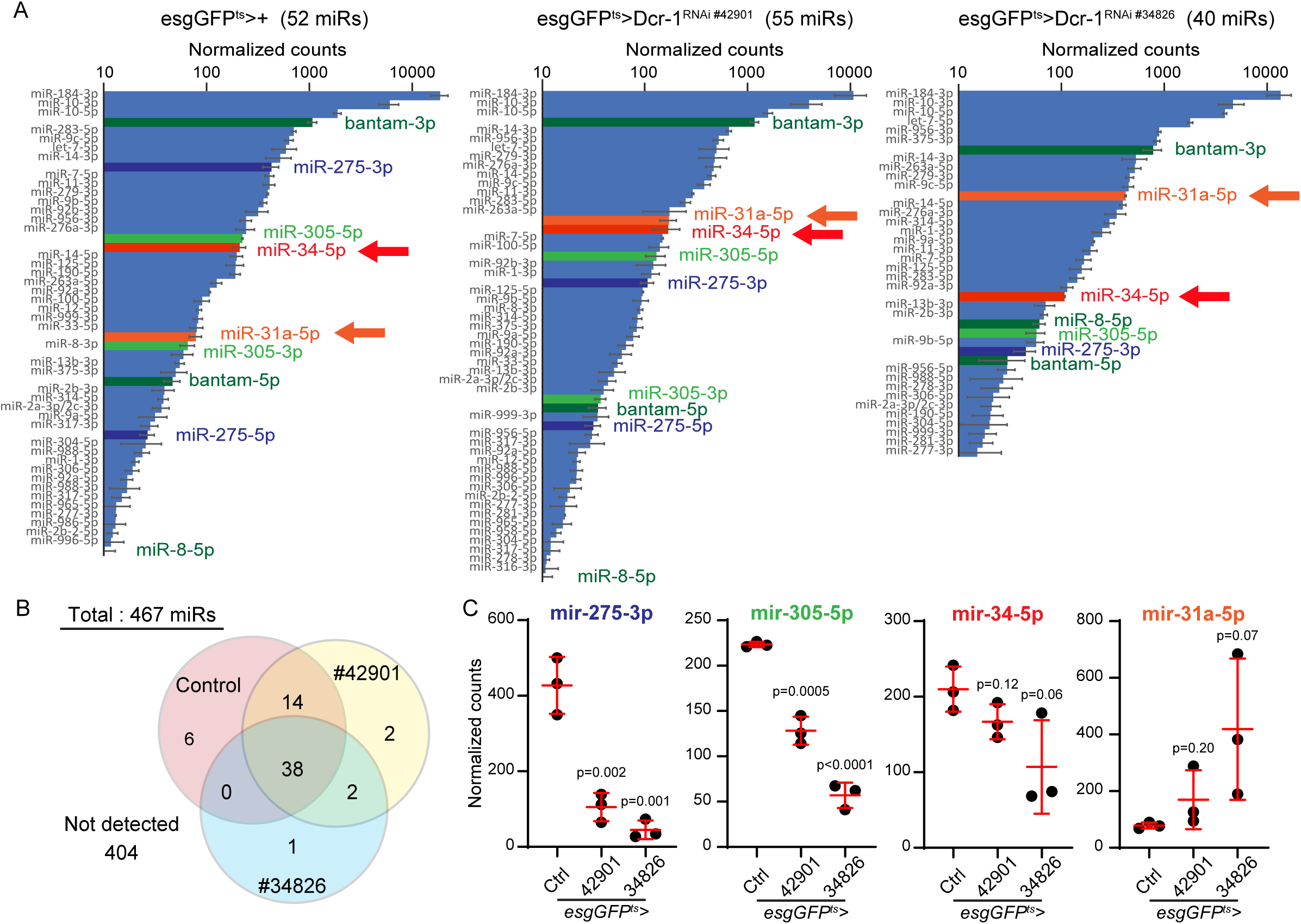
Detecting microRNAs in isolated adult esg-positive intestinal cells. A. Ranked list of the microRNAs detected in each genetic condition (controls and 2 esg-specific Dcr-1 knock downs). miRs detected with a normalized count of more than 5 in each replicate and an average of more than 10 counts across triplicated are shown. B. Venn diagram presenting the number of miRs detected (>5 normalized counts in each biological replicate for a specific genetic conditions). 38 miRs were detected in all libraries. C. Specific list of microRNAs detected in each genetic conditions (control and 2 esg-specific Dcr-1 knockdowns) which have been previously identified to play role in intestinal stem cells (miR-275 and miR-305) along with 2 microRNAs in this study (miRs detected with a normalized count of more than 5 in each replicate.

Remarkably, our strategy allowed us to detect the expression of all the miRs that have previously been described in the intestinal lineage (bantam, miR-275, miR-305 and miR-8): (i) bantam (Huang, Li et al. 2014); (ii) mir-275 (Foronda, Weng et al. 2014); (iii) mir-305 (Foronda, Weng et al. 2014); (iv) mir-8 controls late stage EB differentiation and the expression of *escargot* (Antonello, Reiff et al. 2015). For miRs that have no known function in the intestinal lineage, we detected strand specific biases (e.g. mir-31a-5p and mir-34-5p, but not mir-31a-3p and mir-34-3p) similar to the biases described in previous microRNA sequencing studies from brain and other tissues ({Liu, 2012 #25}{Xiong, 2016 #30}{Srinivasan, 2022 #24}{Foo, 2017 #21}{Jung, 2020 #20}; collectively described in miRbase. Altogether, these results confirm the specificity of our approach and the reliability of the miR sequences detected from sorted esg-positive cells.

Of note, detected miRs respond differently to Dcr-1 knock down. For example, the expression of mir-9, mir-275 and mir-305 is significantly decreased in esg>Dcr-1^RNAi^ esg-positive cells, while miR-1 levels are significantly elevated in Dcr-1 loss-of-function cells (Figure 2D and S1D). Based on the relative absence of EBs among esg-positive cells in Dcr-1 knock-down conditions, this suggests that miRs that showed increased representation in these samples are enriched in ISCs compared to EBs (e.g. miR-1). Conversely, miRs that show reduced expression in Dcr-1^RNAi^ esg-positive cells are likely enriched in EBs (e.g. mir-9, mir-275 and mir-305). Alternatively, distinct miRs may employ different biosynthetic pathways or respond to different compensatory signals in response to Dcr-1 loss-of-function.

**Figure 2:**
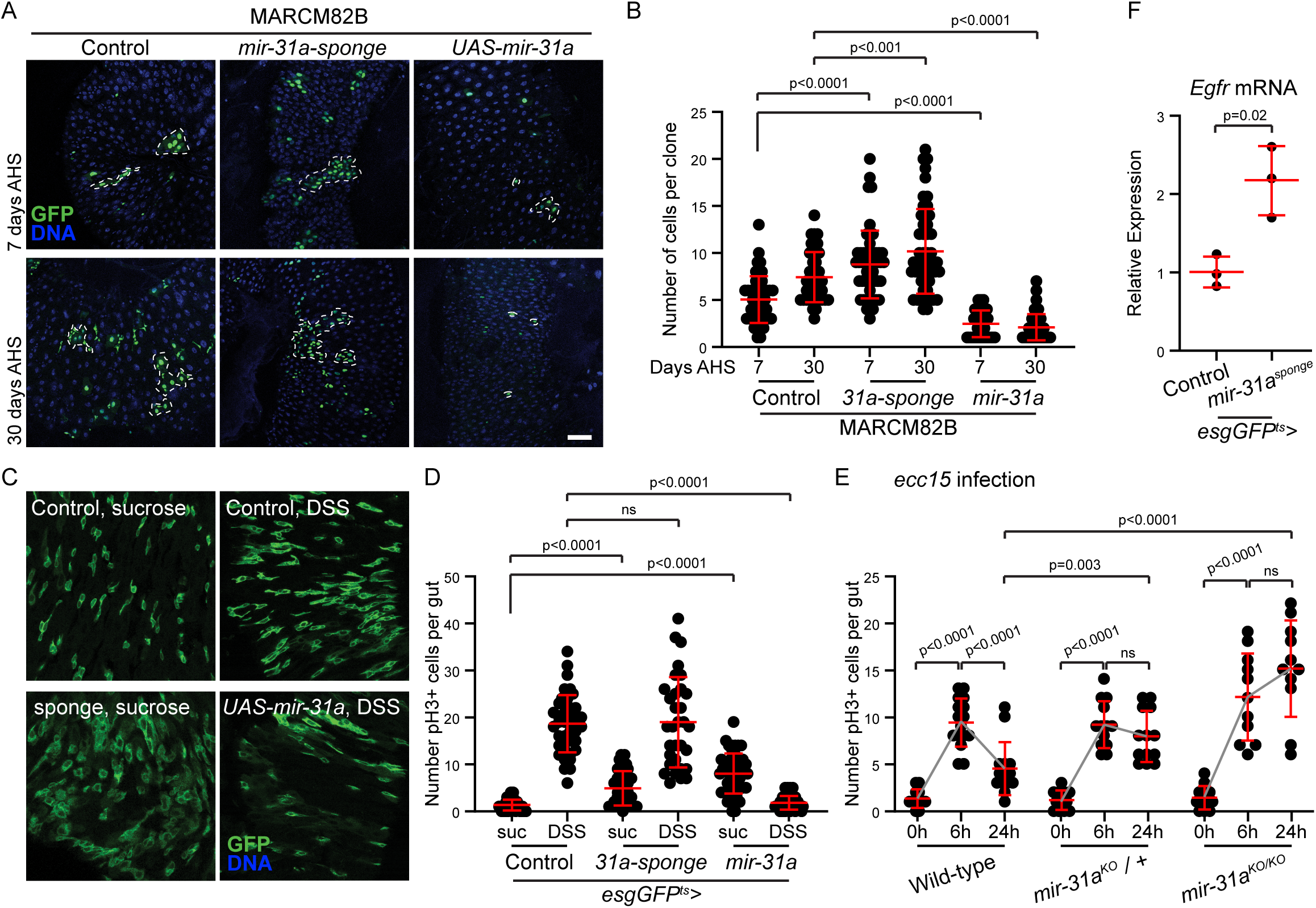
miR-31a negatively controls ISC proliferation and is required for their re-entry into quiescence. A,B. MARCM analysis demonstrating the impact of miR-31 loss-and gain-of-function on ISC proliferation. GFP-positive were imaged 7 and 30 days after induction by heat shock. Clone size for all conditions is measured in B. C,D represent the effect of DSS treatment on various genetic conditions of conditional loss and gain-of function of miR31 using miR31 sponge and UAS-miR31a. The number of pH3+ve cells were counted for each condition. Each data point represents an individual gut in D. E represents the effect of pathogenic (ecc15) insult on miR31 mutant. The proliferative In panels A and C, DNA stained using Hoechst is presented in blue and GFP is in green. In panels B, D and E, individual data points are represented as well as Average +/-standard deviation; p-values are calculated using Student’s t-test.

To improve our understanding of the critical microRNAs involved in the regulation of the intestinal epithelium lineage, we genetically manipulated the activity of several conserved miRs detected with high confidence. Of those, mir-31a and mir-34 were selected for further analysis, based on their conservation across organisms and lineages, and on their strong effect on ISC biology.

### mir-31a negatively controls ISC proliferation and their re-entry into quiescence

In *Drosophila*, mir-31a has recently been found to affect wing development and maintain glial homeostasis in the brain (Foo, Song et al. 2017, Jung, Lee et al. 2020). In mammals, numerous and opposite functions of microRNA-31 have been established during development and tumor formation (Stepicheva and Song 2016, Yu, Ma et al. 2018).

To start addressing the function of mir-31a in the ISC lineage, we performed MARCM clonal analysis using UAS-driven specific sponge (loss-of-function) and over-expression (gain-of-function) constructs. As previously shown, control clones progressively grow to an average size of ∼8 cells/clones (Figure 2A). Under the same standard conditions, inhibition of miR-31a leads to a significant accelerated and elevated clone growth. Conversely, increased mir-31a expression severely impairs clone growth. To support these results, we next tested the impact of mir-31 on the ability of ISCs to respond to tissue damage (Figure 2B). In controls animals, exposure to DSS (Dextran Sulfate Sodium) triggers a solid proliferative response in the intestinal epithelium, as measure by the increase in pH3+ (phospho-Histone H3 positive) cells and an expansion of the esg-positive cell population in the midgut. Mirroring the results of our lineage analysis, mir-31 over-expression is sufficient to entirely block this response, supporting the notion that it strongly and negatively affects ISC proliferation.

Interestingly, under these stress conditions, expression of the mir-31 sponge in esg-positive cells does not significantly change the proliferative response (Figure 2B). This absence of loss-of-function phenotype raised the possibility that mir-31a may affect the dynamics of stress response rather than the maximal ISC proliferation following DSS feeding. For example, we previously reported that Tis11 is required for the re-entry of ISCs into quiescence, and that Tis11 loss-of-function results in larger clones but no change in DSS response (McClelland, Jasper et al. 2017). To test this hypothesis and alleviate possible caveats associated with the use of a sponge, we exposed mir-31a null animals to the mild pathogenic bacteria ecc15. As opposed to DSS, ecc15 infection results in a moderate and transient response in the intestine of control animals: proliferation rates reach peak around 6 hours after feeding of a bacteria-laced solution and return to basal levels by 24 hours after exposure (Figure 2C). We found that, in animals heterozygous or homozygous for a mir-31a allele, the initial proliferation (6 hours timepoint) is similar to controls. However, in heterozygous or homozygous animals’ proliferation remains high 24 hours after infection.

Altogether these data demonstrate that mir-31a is a negative regulator of ISC proliferation and suggest that it contributes to the regulation of the re-entry of ISCs into quiescence after episodes of tissue turnover. High confidence mir-31a predicted targets include the Egfr (Epidermal Growth Factor Receptor) Table S1 {Agarwal, 2018 #35}. These are known components of signaling pathways required for ISC proliferation ({Jin, 2015 #34}), strongly suggesting that mir-31a regulates ISC by controlling at least some of these messengers. Thus, we used RT-qPCR to measure the expression levels of the *Egfr* in mir-31a mutants. Fig 2F

### miR-34 levels strongly impact ISC cell division cell autonomously

The mir-34 family is highly conserved from arthropods to vertebrates (Wang, Jia et al. 2022). In *Drosophila*, mir-34 has been mostly investigated in the developing and aging brain and the immune system (Liu, Landreh et al. 2012, Xiong, Kurthkoti et al. 2016, McNeill, Warinner et al. 2020, Srinivasan, Tran et al. 2022). In humans, mir-34 targets a variety of signaling pathways, for example to control epithelial-to-mesenchymal transition and apoptosis in diverse cancer types (Misso, Di Martino et al. 2014, Zhang, Liao et al. 2019).

Mirroring our analysis of mir-31a, we first performed MARCM clonal analysis to test the impact of mir-34 null mutation on ISC lineages. We found that homozygous null clones remain as single GFP-positive cells, as compared to control clones that grow to an average of 7-8 cells, 7 days after induction (Figure 3A and 3B). Interestingly, mir-34 null animals are viable; thus, we compared their intestinal cell composition to controls animals. Supporting the notion that loss of mir-34 results in decreased ISC proliferation under homeostatic conditions, we found that EBs (Sox21a-positive Delta-negative cells) are almost absent from the intestine of mir-34 null homozygotes, despite a slight increase in the proportion of ISCs (Sox21a and Delta double positive cells) (Figure 3C and D). Next, we tested the ability of mir-34 mutant ISCs to respond to tissue damage. DSS treatment induces a robust proliferative response in the intestine of control animals (Figure 3E). However, this response is significantly diminished in mir-34 heterozygotes and absent in mir-34 null mutants.

**Figure 3:**
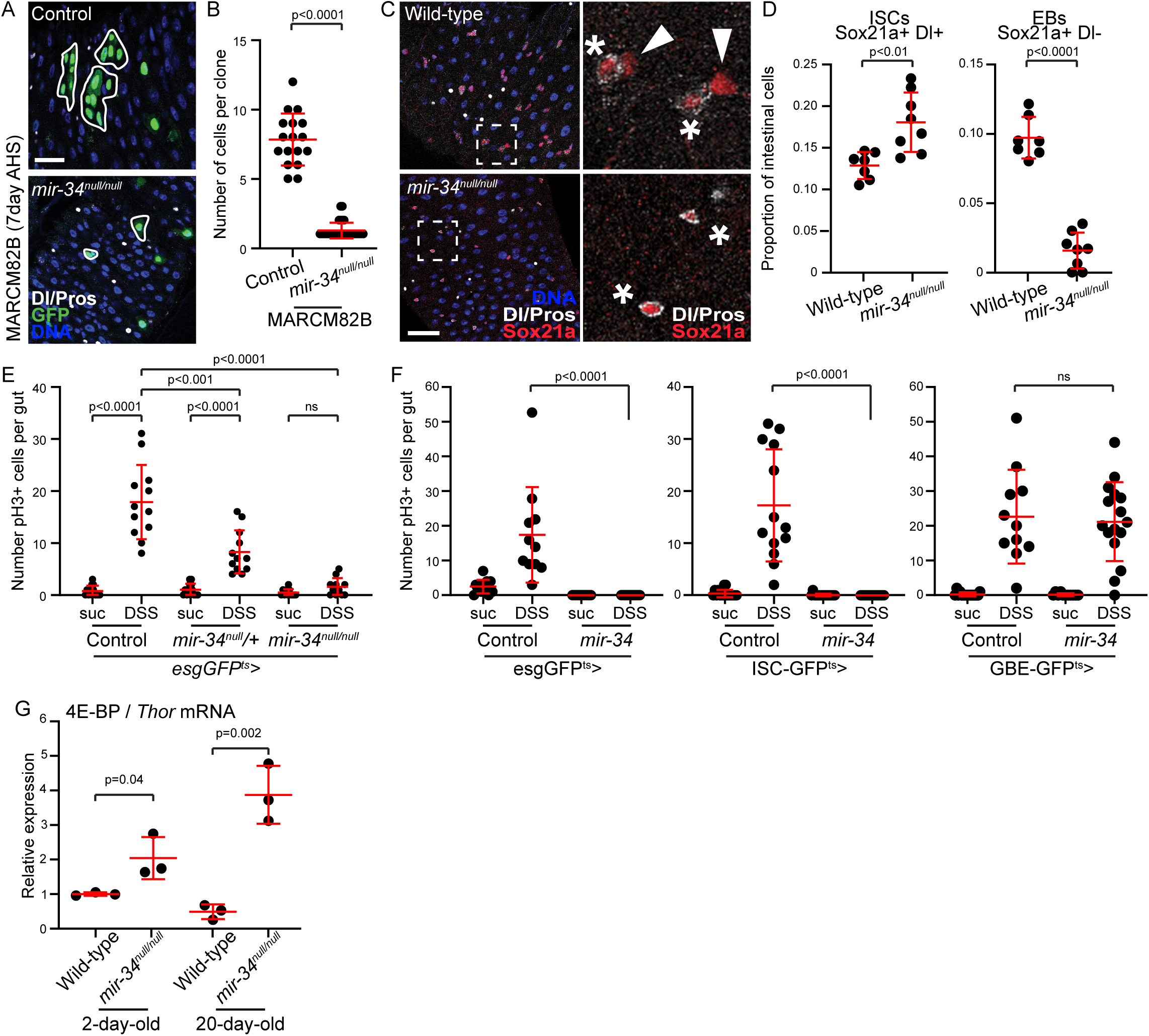
miR-34 is required for ISC proliferation. A.B MARCM clonal analysis of miR34 mutants shows block in ISC proliferation 7 days post clonal induction as compared to control animals. B represents the number of cells per clone. C,D Wildtype and miR34 mutant guts stained with ISC marker delta and ISC-EB marker Sox21a. D represents the quantification of number of ISC and EBs in the wildtype and miR34 guts. E,F demonstrates the effect of DSS treatment on miR34 guts. DSS treatment does not induce proliferation in miR34 mutants, whereas cell specific expression of miR34 suggests that miR34 is required in ISC’s not EB’s. G shows the relative expression of Thor mRNA in the guts in an age dependent manner. In panels A, DNA is stained using Hoechst and GFP is green, in panel C, ISC’s are stained using Delta in white and ISC’s and EB’s are stained together using Sox21a in red. P-values are calculated using student’s t-test.

Next, we sought to investigate the impact of mir-34 gain-of-function in the intestinal lineage. We drove the expression of mir-34 in ISCs and EBs using esgGFP^ts^ driver, only in ISCs using the ISC-GFP^ts^ driver and in EBs using the GBE-GFP^ts^ driver. While intestinal proliferation is robustly induced by DSS in all control conditions, ISC proliferation is abolished when mir-34 is over-expressed in ISCs but not significantly affected when over-repression is limited to EBs (Figure 3F).

Target prediction analysis as described earlier (Table S1, Ref) shows Thor(4E-BP) as one of the targets, where Thor expression is associated with regulation of cell growth and metabolism in *Drosophila* midgut(Ref). To this end, we used RT-qPCR to measure the relative expression of *thor* in an age dependent manner. Interestingly loss of miR-34 in the gut leads to higher expression of *thor*-mRNA suggesting that miR-34 potentially regulates Thor (Fig. 3G).

Altogether, our sequencing approach and validation of the microRNAs in the Intestinal progenitor would be useful for the functional characterization of microRNAs. Further studies are required to determine the role of these microRNAs stem cell proliferation, growth and metabolism.

## Materials and methods

### Fly stocks and husbandry

The following stocks were obtained from Bloomington Drosophila Stock Center; *w^1118^*, *UAS-Dcr1* RNAi (*y1 sc* v1 sev21; P{TRiP.HMS02594}attP40*; 42901, (*y1 sc* v1 sev21; P{TRiP.HMS00141}attP2*; 34826), *neoFRT82B*, *UAS-mCherry-miR-31a-Sponge*(*w*; P{UAS-mCherry.mir-31a.sponge.V2}attP40; P{UAS-mCherry.mir-31a.sponge.V2}attP2*; 61383), *UAS-miR-31a* (*w1118; P{UAS-LUC-mir-31a.T}attP2*; 42027), *miR-31a-KO*( *w*; TI{TI}mir-31aKO*; 58928), *miR-34 ^null/null^* (a gift from Nancy M Bonini), MARCM82B (*hsflp,UAS-GFP;tub-gal4; FRT82B tub-Gal80*) was a gift from N. Perrimon. Intestinal progenitor-specific driver *esgGal4,UAS-GFP;tub-Gal80^ts^* (termed *esgGFP^ts^*) throughout this manuscript. *neoFRT82B.UAS-mir31a*, *neoFRT82B.UAS-mCherry.miR31a.sponge*, *neoFRT82B.miR34^null/null^* (this study). The flies were reared in standard cornmeal/agar medium supplemented with yeast at 25°C with 60±5% relative humidity and 12h light /dark cycles unless specified.

### Conditional Expression of UAS-Linked transgenes

*tub-Gal80^ts^* was used to suppress the early activity of Gal4 before adulthood by rearing them at 18°C in *esgGFP^ts^*. 2–3-day old adults were transferred to 29°C to activate Gal4. The flies were reared at 29°C in standard food unless specified. Gal4 drivers crossed to *w^1118^* were used as control for these experiments.

### Mosaic analysis with repressible cell marker (MARCM) clones

4-5 days old flies were heat shocked 3 times at 37°C for 45min to 1 hour within one day for MARCM analysis. The files were maintained at 25°C unless specified. MARCM82B were used for inducing somatic clones. For analysis of the induction of GFP positive clones, appropriately aged 6-7 female guts for each genotype were quantified for the number of cells per clone in the posterior midgut region.

### Stress exposure and infection

Dextran Sodium Sulphate (DSS, 4%; Sigma-Aldrich) and Ecc15 infection were used for in vivo stress experiments. For DSS treatment, young adult flies were starved for 4hours followed by feeding them with 4% DSS in 5% sucrose-saturated filter paper. Flies fed with 5% sucrose only were used as controls. Intestines were dissected 36 hours after treatment.

For *Ecc15* (*Erwinia carotovora carotovora 15*) treatment, pellet from the overnight culture of *Ecc15* (OD_600_=200) was mixed in 5% sucrose and added to the vials containing flies starved for 4hours. Intestines were dissected at specific times as indicated in figure legends. Flies fed with 5% sucrose only were used as controls.

### Immunohistochemistry

Guts were immunostained as described previously. Guts were dissected in ice cold PBS and immediately fixed in glutamate buffer containing 4% formaldehyde for 20 minutes with equal volume of heptane. Following fixation, the guts were washed in methanol (100%, 70%, 50% in 1XPBS and final wash of 25% methanol in 1XPBS with 0.1% Tween-20). Following methanol washes the guts were washed in PBS with 0.1% TX-100 and 0.5% BSA (gut buffer). The guts for blocked in the same buffer for one hour and incubated with primary antibody overnight at 4°C. Secondary antibody was used at a dilution of 1:500. The guts were finally washed in gut buffer and mounted on slides with Mowiol/Dabco solution. The following antibodies were used: Sox21a (1:50000; previously generated in the lab); anti-Delta (1:500, C594.9B), anti-Prospero (1:500, MR1A) were obtained from Developmental Studies Hybridoma Bank (DSHB) and Anti-pH3 (1:2000, 06-570) from Millipore. Fluorescent secondary antibodies were obtained from Jackson Immunoresearch. Hoechst 33258 (Sigma Aldrich) was used to stain DNA.

### FACS sorting of intestinal progenitors

Appropriately aged guts of specific genotypes (approx. 100 guts per genotype) were dissected in ice cold 1XPBS (DEPC) according to protocol as described previously (Ref). *w^1118^* was used as negative control. The guts were digested with Collagenase-IV (10mg/ml in 1XPBS-DEPC). Incubate the guts at 27°C for 1 hour with slow agitation. The guts were agitated every 15 min with pipette for 30-40 times for full dissociation. The dissociated cells were centrifuges at 300rpm at 4°C for 20 mins. The pellet containing the cells were resuspended in ice cold PBS-DEPC and processed for FACS analysis with 1 µl of propidium iodide. The cells were sorted based on GFP fluorescence and size using the BD FASC Aria II system.

### Small RNA seq analysis of FACS sorted cells

Total RNA was extracted from sorted GFP positive RFP negative cells using *mir*Vana^TM^ miRNA Isolation kit (Invitrogen) according to manufactures protocol. miRNA seq analysis was performed at UR Genomics Research center. The total RNA concentration was determined with Nanodrop 1000 spectrophotometer (Nanodrop, Wilmington, DE) and RNA quality assessed with the Agilent Bioanalyzer (Agilent, Santa Clara, CA). 500 ng of Total RNA was utilized as the input for sequencing library construction with the NEB Next small RNA library prep kit (New England Biolabs), following the manufacturer’s protocol. Importantly, 3’ and 5’ adapters were diluted 2-fold for ligation steps, as specified by the manufacturer. Following 3’adaptor ligation, RT primer incubation, and 5’ adapter ligation, cDNA was synthesized, and final library amplification was performed with 12 cycles of PCR. Library quantity and quality were assessed using a Qubit fluorometer (Thermofisher) and Tapestation 2200 (Agilent), respectively. Sample libraries were proportionally pooled and isolated for the PCR reaction mix using the Qiagen MinElute PCR Reaction Cleanup kit and Library fragments between 125-160bp were size selected using a 3% cassette on a PippinHT (Sage Science). The size selected library pool was then sequenced on an illumina HiSeq 2500, generating single end reads of 50nt.

### Demultiplexing, Alignment, and Analysis

Raw reads generated from the Illumina base calls were demultiplexed using bcl2fastq version 2.19.0.miRge2.0 (2.0.3) was used to align reads with bowtie (1.2.1.1) (Langmead, Trapnell et al. 2009) and identify microRNAs with the following project specific parameter “-sp. Fruitfly’. (Ambros, Bartel et al. 2003)

### qRT-PCR analysis

RNA isolated from the FACS sorted cells were used to synthesize cDNA with 50µg of total RNA using Superscript-III reverse transcriptase (Invitrogen). Real-time PCR was performed using the Applied biosystems Quanta Studio using Quantabio Perfecta SYBR green mix according to manufacturer’s protocol using following primers:

*Delta* Forward 5’ CACCTGCGATCTCAACTACTAC 3’.

*Delta* Reverse 5’ GCCATCCGGTCAAACAGATA 3’.

*GFP* Forward 5’ TCAAGATCCGCCACAACATC 3’.

*GFP* Reverse 5’ GTGCTCAGGTAGTGGTTGTC 3’.

*Escargot* Forward 5’ CCGCCCATGAGATCTGAAAT 3’.

*Escargot* Reverse 5’ GGTGATGATGGGTATGGGTATAG 3’.

*rp49* Forward 5’ CCAGTCGGATCGATATGCTAAG 3’.

*rp49* Reverse 5’ CCGATGTTGGGCATCAGATA 3’.

*Thor* Forward 5’ATGCAGCAACTGCCAAATC 3’.

*Thor* Reverse 5’ GAGAACAAACAAGGTGGAAGAAC 3’.

*EGFR* forward 5’ CAAGAGCAGGGATCGCTAAA 3’.

*EGFR* Reverse 5’ CACCTGTTCATGGTATCCGTAG 3’.

Relative expression was calculated using the ΔΔCT method and normalized to rp49 levels. P values were calculated using unpaired two-tailed student’s t-test.

### Image analysis

Confocal images were collected using a Leica SP5 confocal system and processed using the Leica-LAS-X software and analyzed using the Fiji/Image J software and assembled using Adobe illustrator.

### Statistics

All p-values were calculated using the student’s t-test with unpaired samples. All error bars represent standard error of mean. Exact values of all n’s can be found in Figure legends.

---

**Figure S1.**
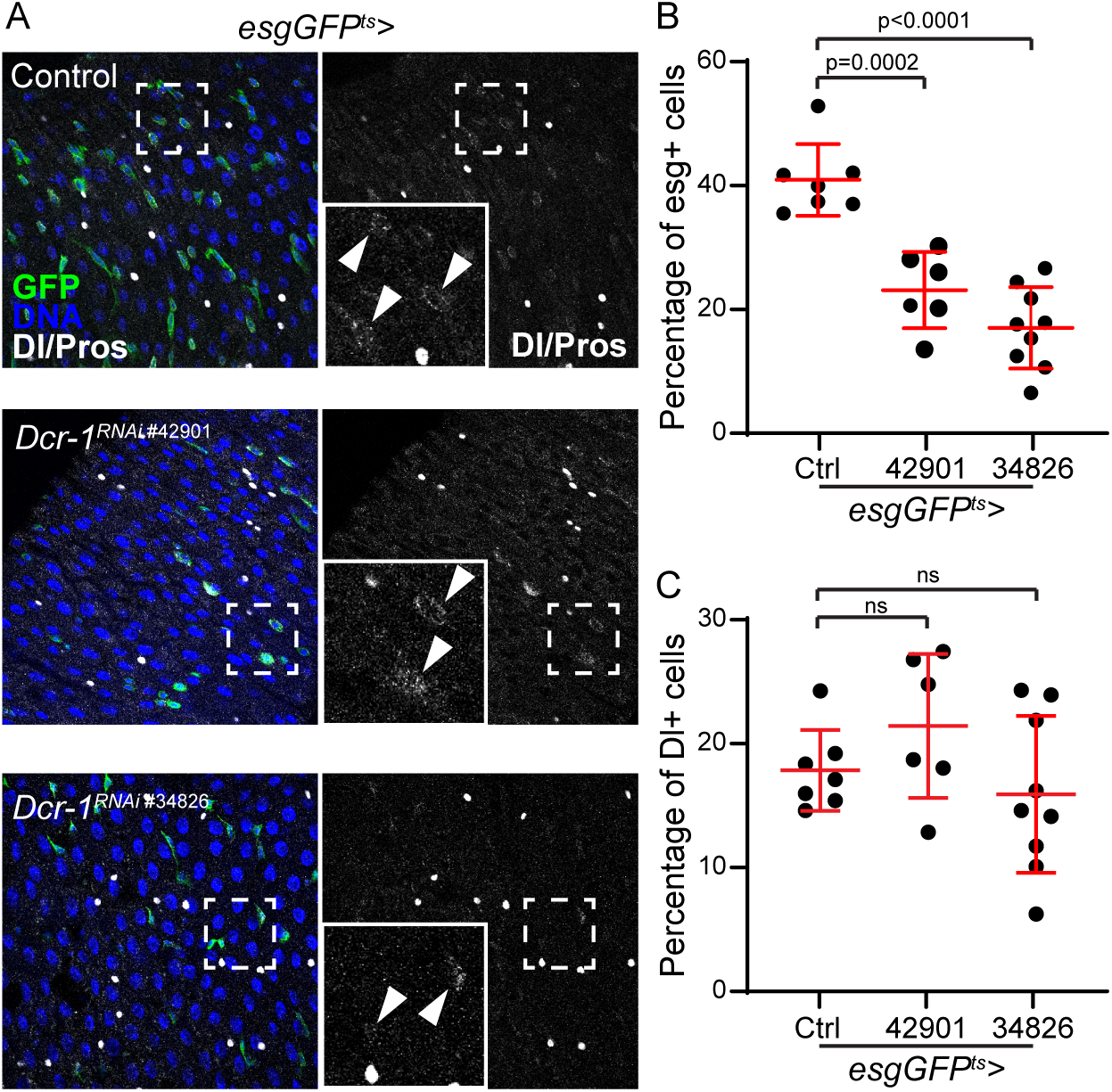

**Figure S2.**
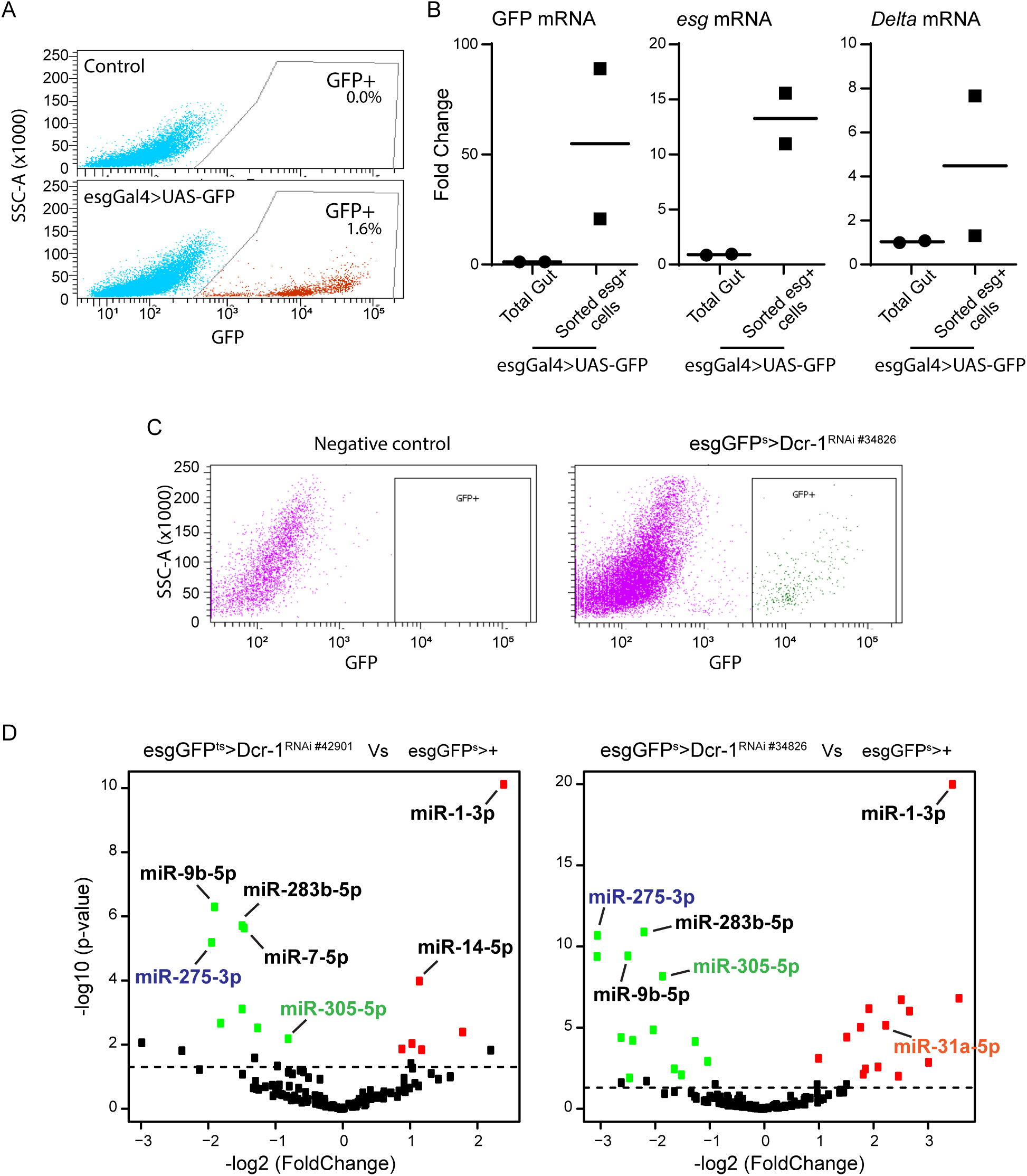

## References

Ambros, V., et al. (2003). "A uniform system for microRNA annotation." RNA 9(3): 277–279.

Antonello, Z. A., et al. (2015). "Robust intestinal homeostasis relies on cellular plasticity in enteroblasts mediated by miR-8-Escargot switch." Embo Journal 34(15): 2025–2041.

Bartel, D. P. (2004). "MicroRNAs: genomics, biogenesis, mechanism, and function." Cell 116(2): 281–297.

Dutta, D., et al. (2015). "Regional Cell Specific RNA Expression Profiling of FACS Isolated Drosophila Intestinal Cell Populations." Curr Protoc Stem Cell Biol 34: 2F 2 1–2F 2 14.

Foo, L. C., et al. (2017). "miR-31 mutants reveal continuous glial homeostasis in the adult Drosophila brain." EMBO J 36(9): 1215–1226.

Foronda, D., et al. (2014). "Coordination of insulin and Notch pathway activities by microRNA miR-305 mediates adaptive homeostasis in the intestinal stem cells of the Drosophila gut." Genes Dev 28(21): 2421–2431.

Huang, H. L., et al. (2014). "Bantam is essential for Drosophila intestinal stem cell proliferation in response to Hippo signaling." Developmental Biology 385(2): 211–219.

Jiang, H., et al. (2016). "Intestinal stem cell response to injury: lessons from Drosophila." Cell Mol Life Sci 73(17): 3337–3349.

Jonas, S. and E. Izaurralde (2015). "Towards a molecular understanding of microRNA-mediated gene silencing." Nat Rev Genet 16(7): 421–433.

Jung, J. E., et al. (2020). "MicroRNA-31 Regulates Expression of Wntless in Both Drosophila melanogaster and Human Oral Cancer Cells." Int J Mol Sci 21(19).

Kim, K., et al. (2017). "miR-263a Regulates ENaC to Maintain Osmotic and Intestinal Stem Cell Homeostasis in Drosophila." Developmental Cell 40(1): 23–36.

Lam, V., et al. (2014). "bantam miRNA is important for Drosophila blood cell homeostasis and a regulator of proliferation in the hematopoietic progenitor niche." Biochemical and Biophysical Research Communications 453(3): 467–472.

Langmead, B., et al. (2009). "Ultrafast and memory-efficient alignment of short DNA sequences to the human genome." Genome Biol 10(3): R25.

Liu, N., et al. (2012). "The microRNA miR-34 modulates ageing and neurodegeneration in Drosophila." Nature 482(7386): 519–523.

Lu, Y., et al. (2018). "miRge 2.0 for comprehensive analysis of microRNA sequencing data." BMC Bioinformatics 19(1): 275.

McClelland, L., et al. (2017). "Tis11 mediated mRNA decay promotes the reacquisition of Drosophila intestinal stem cell quiescence." Developmental Biology 426(1): 8–16.

McNeill, E. M., et al. (2020). "The conserved microRNA miR-34 regulates synaptogenesis via coordination of distinct mechanisms in presynaptic and postsynaptic cells." Nat Commun 11(1): 1092.

Misso, G., et al. (2014). "Mir-34: a new weapon against cancer?" Mol Ther Nucleic Acids 3(9): e194.

Mukherjee, S., et al. (2021). "A stress-responsive miRNA regulates BMP signaling to maintain tissue homeostasis." Proceedings of the National Academy of Sciences of the United States of America 118(21).

Shcherbata, H. R. (2019). "miRNA functions in stem cells and their niches: lessons from the Drosophila ovary." Current Opinion in Insect Science 31: 29–36.

Singh, A. P., et al. (2020). "Enteroendocrine Progenitor Cell-Enriched miR-7 Regulates Intestinal Epithelial Proliferation in an Xiap-Dependent Manner." Cell Mol Gastroenterol Hepatol 9(3): 447–464.

Srinivasan, A. R., et al. (2022). "Loss of miR-34 in Drosophila dysregulates protein translation and protein turnover in the aging brain." Aging Cell 21(3): e13559.

Stepicheva, N. A. and J. L. Song (2016). "Function and regulation of microRNA-31 in development and disease." Mol Reprod Dev 83(8): 654–674.

Toledano, H., et al. (2012). "The let-7-Imp axis regulates ageing of the Drosophila testis stem-cell niche." Nature 485(7400): 605–610.

Wang, C., et al. (2022). "microRNA-34 family: From mechanism to potential applications." Int J Biochem Cell Biol 144: 106168.

Wu, Y. C., et al. (2017). "The bantam microRNA acts through Numb to exert cell growth control and feedback regulation of Notch in tumor-forming stem cells in the Drosophila brain." Plos Genetics 13(5).

Xiong, X. P., et al. (2016). "miR-34 Modulates Innate Immunity and Ecdysone Signaling in Drosophila." PLoS Pathog 12(11): e1006034.

Yang, Y. Y., et al. (2009). "The Bantam microRNA Is Associated with Drosophila Fragile X Mental Retardation Protein and Regulates the Fate of Germline Stem Cells." Plos Genetics 5(4).

Yu, T., et al. (2018). "Functions and mechanisms of microRNA-31 in human cancers." Biomed Pharmacother 108: 1162–1169.

Zhang, L., et al. (2019). "MicroRNA-34 family: a potential tumor suppressor and therapeutic candidate in cancer." J Exp Clin Cancer Res 38(1): 53.

Zipper, L., et al. (2022). "The MicroRNA miR-277 Controls Physiology and Pathology of the Adult Drosophila Midgut by Regulating the Expression of Fatty Acid beta-Oxidation-Related Genes in Intestinal Stem Cells." Metabolites 12(4).

